# Cognitive limits of larval *Drosophila*: Testing for conditioned inhibition, sensory preconditioning and second-order conditioning

**DOI:** 10.1101/2023.08.21.554112

**Authors:** Edanur Sen, Amira El-Keredy, Nina Jacob, Nino Mancini, Gülüm Asnaz, Annekathrin Widmann, Bertram Gerber, Juliane Thoener

## Abstract

*Drosophila* larvae are an established model system for studying the mechanisms of innate and simple forms of learned behaviour. They have about 10 times fewer neurons than adult flies, and it was the low total number of their neurons that allowed for an electron microscopic reconstruction of their brain at synaptic resolution. Regarding the mushroom body, a central brain structure for associative learning in insects, it turned out that more than half of the classes of synaptic connection had previously escaped attention. Understanding the function of these circuit motifs, subsequently confirmed in adult flies, is an important current research topic. In this context, we test larval *Drosophila* for their cognitive abilities in three tasks that are characteristically more complex than those previously studied. Our data provide evidence for (i) conditioned inhibition, as has previously been reported for adult flies and honeybees. Unlike what is described for adult flies and honeybees, however, our data do not provide evidence for (ii) sensory preconditioning or (iii) second-order conditioning in *Drosophila* larvae. We discuss methodological features of our experiments as well as four specific aspects of the organisation of the larval brain that may explain why these two forms of learning are observed in honeybees and adult flies, but not in larval *Drosophila*.

## Introduction

Investigations of *Drosophila melanogaster* have led to the discovery of evolutionarily conserved mechanisms of development and the function of ion channels and synapses, and shed light on how synaptic plasticity and circadian rhythms are neurogenetically organized (Johnston and Nüsslein-Volhard 1992; Littleton and Ganetzky 2000; Foster and Helfrich-Förster 2001). Notably, conserved mechanisms were also discovered for associative learning and memory (Dudai et al. 1976; Heisenberg et al. 1985; Tully and Quinn 1985). These analyses gained momentum when combined with convenient methods for cell-specific transgene expression (Brand and Perrimon 1993; Pfeiffer et al. 2010) to study the neuronal circuits underlying this simple form of cognition (Zars et al. 2000; Guven-Ozkan and Davis 2014; Gerber and Aso 2017; Cognigni et al. 2018; Boto et al. 2020).

The potential of larval *Drosophila* for the analysis of learning and memory was realized early on (Aceves-Piña and Quinn 1979) and with renewed interest when paradigms were established for the association of odours and taste reward (Scherer et al. 2003; Neuser et al. 2005) and between odours and the optogenetic activation of brain reward neurons (Schroll et al. 2006). Larvae possess about 10 times fewer neurons than adult flies, but feature adult-like circuit motifs, for example in the olfactory pathways (Gerber and Stocker 2007; Vosshall and Stocker 2007; Diegelmann et al. 2013; Thum and Gerber 2019; Eschbach and Zlatic 2020). The larva’s nervous system has been partially mapped into a light-microscopic cell atlas (Li et al. 2014), and transgenic driver strains can be generated to manipulate these cells, individually or in small groups, in the context of learning experiments (Rohwedder et al. 2016; Saumweber et al. 2018). Furthermore, the electron microscopic reconstruction of the chemical-synapse connectome of a larval brain has revealed unexpected complexity in the mushroom body (Eichler et al. 2017; Winding et al. 2023), a central brain structure for associative learning in larvae and in insects in general. Indeed, more than half of the classes of connection had escaped earlier attention! The functional analysis and computational interpretation of these circuit motifs, subsequently confirmed in adult flies (Takemura et al. 2017; Zheng et al. 2018; Li et al. 2020), now largely define the research agenda of the field. One element of this undertaking is to probe the cognitive limits associated with a mushroom body that is numerically as simplified as in larval *Drosophila*. In this context we test these animals in three tasks, well established in contemporary experimental psychology (Rescorla 1988a, 1988b), that are characteristically more complex than the simple paradigms hitherto applied in larvae: conditioned inhibition, sensory preconditioning, and second-order conditioning.

### Conditioned inhibition

In a typical associative learning task, a cue (A) is presented together with, for example, a reward (+). Such ‘paired’ training (A+) allows the cue to guide learned behaviour in a later test. This is often called ‘conditioned excitation’ because the cue is said to *excite* the expectation of the reward to occur, and to prompt behaviour *in anticipation of receiving* it. In contrast, ‘conditioned inhibition’ refers to the opposing process, which allows the cue to *inhibit* the expectation of the reward to occur, and to prompt behaviour *in anticipation of not receiving* it (Rescorla 1969). Conditioned inhibition can be established, for example, by training the subjects such that precisely whenever the reward is presented the cue is not presented, and vice versa (‘unpaired’ training: +/A). In other words, whereas paired training establishes A as a predictor of reward occurrence (conditioned excitation), unpaired training establishes A as a predictor for the reward’s non-occurrence (conditioned inhibition). Learning through unpaired training is characteristically complex because although it is about the reward, it takes place at a moment when the reward is not physically present.

Opposing effects of paired versus unpaired training have been reported in larval *Drosophila* (Saumweber et al. 2011; Schleyer et al. 2018, 2020), adult flies and honeybees *Apis mellifera* (Bitterman et al. 1983; Matsumoto et al. 2012; Barth et al. 2014; Jacob and Waddell 2020). Here we confirm and extend these results in larval *Drosophila* with respect to hallmark features of conditioned inhibition, using presentations of a sugar taste reward unpaired from an odour cue.

### Sensory preconditioning

Sensory preconditioning (Brogden 1939) refers to the learning that results from two cues A and B, say the visual appearance and the song of a bird, occurring together, and that establishes their combination into a psychological object. In other words, it refers to the chunking of inputs into mnemonic objects according to their past co-occurrence. The characteristic complexity of sensory preconditioning is that it takes place in the absence of reinforcement, and that it allows for pattern completion if one of the cues is not physically present, for example when the song of a robin calls up its visual appearance. Experimentally, sensory preconditioning can be demonstrated in a two-stage experiment: first, in phase (i) AB are presented, then in phase (ii) A+ is trained, and finally a test of B follows. Responding to B during the test would be indicative of sensory preconditioning and would require chained processing during phase (ii) such that by virtue of the AB association cue A calls up B, which is then associated with +, and/or chained processing during the test such that, again by virtue of the AB association, cue B calls up A, which then calls up the A+ association (Molet et al. 2012). Thus, either way these processes hinge on the AB association established in phase (i).

Sensory preconditioning has been shown in adult flies and bees (Müller et al. 2000; Brembs and Heisenberg 2001; Martinez-Cervantes et al. 2022), including for binary odour compounds as cues A and B. We sought to establish sensory preconditioning in larvae, likewise between the elements of binary odour compounds.

### Second-order conditioning

Second-order conditioning (Pavlov 1927; Rescorla 1980) refers to the observation that when a cue A is firmly associated with, for example, a reward it can itself have a rewarding effect – even when the reward is not physically present. In humans, money is an example of a second-order reward. Experimentally, second-order conditioning can be demonstrated by first (i) training A+, then (ii) presenting AB, followed by a test of B revealing a response. The conceptual significance of second-order conditioning is that it allows temporally distant goals to be pursued in a chained manner, such as working for money to buy food. Also, for cues repeatedly experienced in succession before a reward, for example O-then-X-then-A-then-reward, second-order conditioning allows the staggered formation of associations for cues such as O, which would be temporally too far removed from the reward if they were trained in isolation. In other words, second-order conditioning is a process that can identify the earliest reward-predicting cue.

Second-order conditioning has been observed in adult flies and bees (Takeda 1961; Bitterman et al. 1983; Brembs and Heisenberg 2001; Hussaini et al. 2007; Tabone and De Belle 2011; Yamada et al. 2023), again including for binary odour compounds as cues. We sought to establish second-order conditioning for larvae, using binary odour compounds as well.

## Materials and Methods

### Animals, materials and chemicals

*Drosophila melanogaster* larvae were raised in mass culture on standard cornmeal-molasses food and maintained at 25°C, 60–70% relative humidity and a 12:12 h light/dark cycle. For behavioural experiments, 5-day-old, 3^rd^ instar, feeding-stage, wild-type Canton Special larvae of either prospective sex were used. Cohorts of approximately 30 larvae were collected from the food vials, rinsed in water, collected in a water droplet, and then used for experiments.

For behavioural experiments, Petri dishes of 9 cm diameter (Nr. 82.1472 Sarstedt, Nümbrecht, Germany) were filled with 1 % agarose solution as the substrate (PUR; electrophoresis grade; CAS: 9012-36-6, Roth, Karlsruhe, Germany) or with fructose as the sugar taste reward (+) added to the agarose solution (2 M; purity 99 %; CAS: 57-48-7 Roth, Karlsruhe, Germany). As will be specified in the *Results* section, for odour presentation either custom-made Teflon containers or Petri dish lids equipped with filter papers were used. The Teflon containers were of 5 mm diameter with perforated lids with 5-10 holes, each of approximately 0.5 mm diameter. These were filled with 10 µl of odour solution before the experiment and used for one day. The aforementioned Petri dish lids were equipped with four filter papers; each of these filter papers was loaded with 5 µl of the respective odour solution shortly before each experiment. When two odours were presented in compound, 5 µl of each odour solution was used per filter paper. The filter papers were renewed after each experiment.

As the odour substances, either *n-*amylacetate (AM; CAS: 628-63-7, Merck, Darmstadt, Germany; diluted 1:20 in paraffin oil; CAS: 8042-47-5, AppliChem, Darmstadt, Germany) or 1-octanol was used (1-OCT; CAS: 111-87-5; Merck, Darmstadt, Germany; undiluted). Paraffin is without behavioural significance in larval *Drosophila* (Saumweber et al. 2011).

### Behavioural experiments

Effects of PAIRED versus UNPAIRED training with one training trial Experiments followed standard procedures, using a one-odour, single-training-trial protocol (Saumweber et al. 2011; Weiglein et al. 2019; for a manual see Michels et al. 2017) (**Fig. 1A**). Larvae underwent either paired or unpaired presentations of AM as the odour and the sugar taste reward (+), followed by a preference test for the odour.

**Figure 1.**
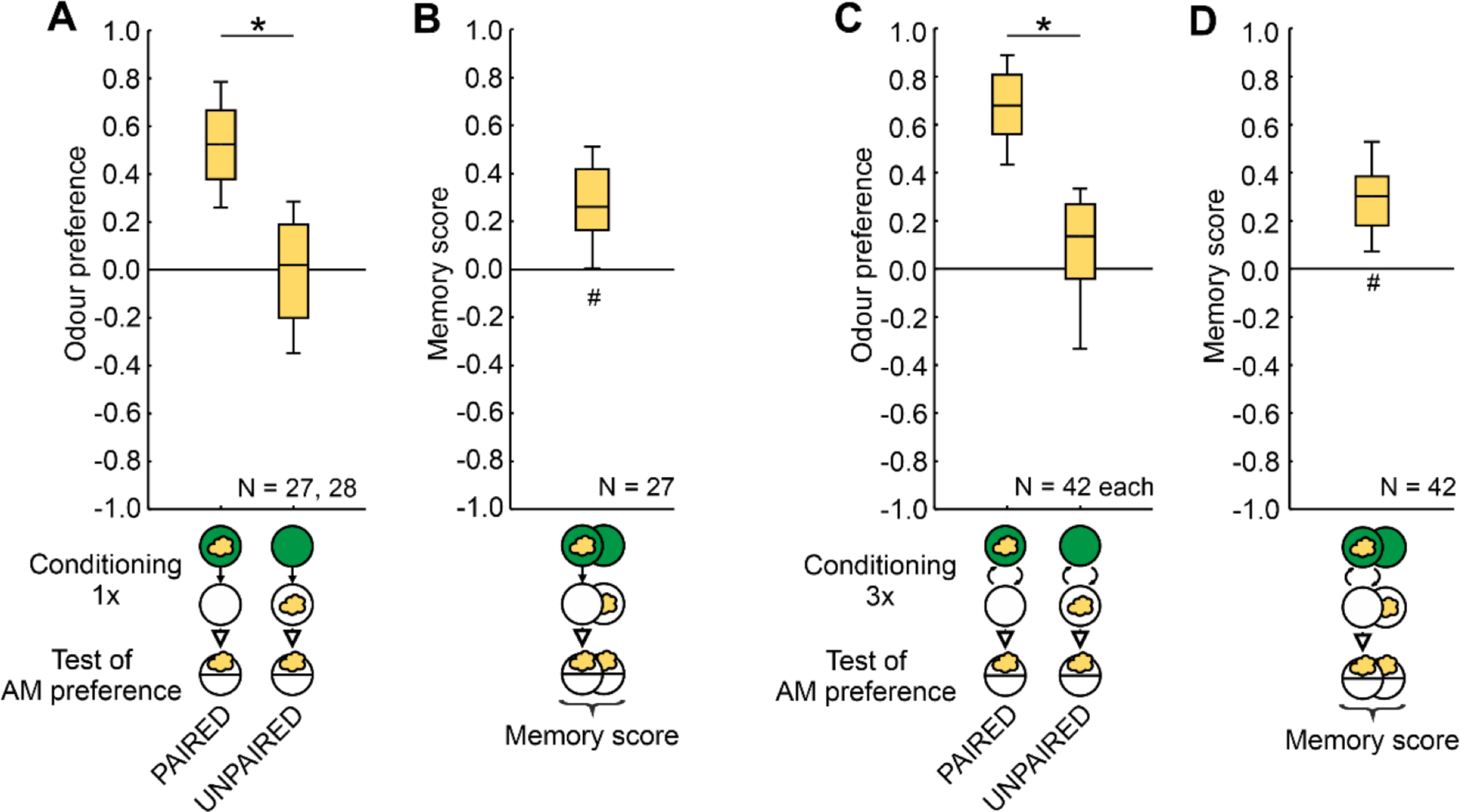
Odour-sugar associative memory in larval *Drosophila*. **(A-D)** *Drosophila melanogaster* larvae received either PAIRED training of the odour (yellow cloud, n-amylacetate) and a fructose sugar reward presented on the same Petri dish (green circle), or they received the sugar reward separately (UNPAIRED training). To equate for training duration and handling, blank periods (open circle) were added for the PAIRED case. The sequence of events during training was as depicted in half of the cases and the reverse in the other half of the cases (not shown). After training, the larvae were tested for their odour preference (Equation 1), and a memory score was calculated from the difference between PAIRED and UNPAIRED training (Equation 2) such that positive and negative memory scores represent appetitive and aversive associative memories, respectively. **(A)** After one-trial PAIRED training the larvae show higher odour preferences than after UNPAIRED training. **(B)** The resulting memory scores indicate appetitive associative memory. **(C, D)** Corresponding results are observed after three training trials. Box plots show the median as the midline, the 25/75% quantiles as box boundaries, and 10/90% quantiles as whiskers. Sample sizes are indicated within the figure. # indicates significance from zero in an OSS test, * indicates significance in an MWU-test. Statistical results and source data in **Supplemental_Table_S1.xlsx**.

For paired training (PAIRED), two containers with AM were located on opposite sides of a Petri dish filled with sugar-supplemented agarose. Cohorts of approximately 30 larvae were placed in the middle of the Petri dish and left undisturbed for 2.5 min to disperse through the Petri dish (AM+). Subsequently, the larvae were collected with a brush and transferred to a second ‘blank’ Petri dish containing neither the sugar reward nor the odour and left free to move about the Petri dish for a further 2.5 min; in this case two empty odour containers (EM) were placed on the Petri dish. In half of the cases this sequence was as mentioned (AM+/EM), whereas in the other half of the cases it was the reverse (EM/AM+). Subsequently, the larvae were transferred to a test Petri dish with agarose but no sugar added (unless mentioned otherwise); in this case an odour container with AM was placed on one side, and an EM container was placed on the opposite side of the Petri dish. After 3 min, the number of larvae on each side (@AM and @EM, respectively) and on a 10-mm-wide middle zone was counted, including the larvae crawling up the sidewalls of the Petri dish; larvae crawling up the lid of the Petri dish were excluded (< 5%). From these numbers a preference score for the odour was calculated as:

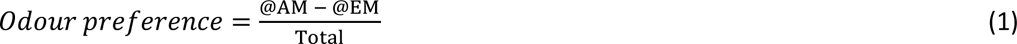

Thus, odour preference scores may range from +1 to -1, with positive values indicating approach to the odour and negative values indicating avoidance.

For unpaired training (UNPAIRED), an independent cohort of larvae was likewise placed in Petri dishes, in this case, however, featuring either only agarose with the sugar reward added but no odour (EM+), or the odour but no sugar reward added to the agarose (AM). Again, either the aforementioned sequence (EM+/AM) or the reverse sequence (AM/EM+) was used, followed by the test for odour preference as described above.

To quantify associative memory, a memory score was calculated as the difference in odour preference after PAIRED versus after UNPAIRED training:

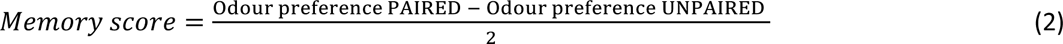

Values for the memory score may thus range from +1 to -1, with positive values indicating appetitive associative memory, and negative values indicating aversive associative memory.

To quantify the effect of using different sequences of odour and sugar reward presentation, we separately calculated a sequence index for the odour preference scores after PAIRED and UNPAIRED training as follows:

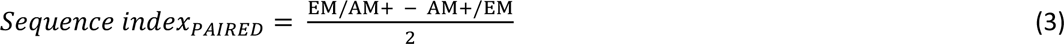

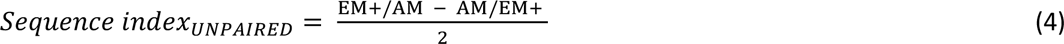

Values of the sequence index may thus range from +1 to -1. In the case of the PAIRED groups, positive values indicate higher odour preferences when the training sequence was EM/AM+ as compared to the reverse Values of the sequence index may thus range from +1 to -1. In the case of the PAIRED groups, positive values indicate higher odour preferences when the training sequence was EM/AM+ as compared to the reverse Values of the sequence index may thus range from +1 to -1. In the case of the PAIRED groups, positive values indicate higher odour preferences when the training sequence was EM/AM+ as compared to the reverse sequence, whereas for the UNPAIRED groups positive values indicate higher preferences when the training sequence was EM+/AM than for the reverse sequence.

In turn, for the PAIRED groups negative sequence indices indicate lower preferences when the training sequence was EM/AM+, whereas for the UNPAIRED groups negative values indicate lower preferences when the training sequence was EM+/AM, as compared to the respective other sequences.

While the arrangement of the numerator and thus the signs of the sequence index are arbitrary in a technical sense, we note that one of the theoretical concepts that we seek to test suggests that conditioned inhibition should lead to negative sequence indices for the UNPAIRED groups if the numerator is arranged as above (see also *Results* section) and that the sequence indices should be zero for the PAIRED groups. We therefore arranged the numerator for the PAIRED groups such that for the data in **Fig. 4D and H** any trend in the sequence indices of the PAIRED group would have the same sign as for the UNPAIRED groups; this is conservative because it underestimates the difference in how strongly the sequence of odour and sugar presentations affects odour preferences in the PAIRED versus the UNPAIRED groups.

#### Effects of PAIRED versus UNPAIRED training with three training trials

All procedures were the same as in the preceding section, except that two additional training trials were performed.

#### Sensory preconditioning

To test for sensory preconditioning, we adapted the paradigm described above. Larvae first underwent a preconditioning phase exposing them to two odours, followed by an odour-reward conditioning phase for one of these odours and a test of preference for the respective other odour (**Fig. 5**). During preconditioning the larvae received the two odours AM and 1-OCT either presented together as a COMPOUND or temporally SEPARATED from each other, in both cases on Petri dishes with agarose but no added sugar; to equate for the total duration of the experiment and for handling we added a blank trial for the COMPOUND case. Three trials of such preconditioning were followed by a conditioning phase, in which the larvae in the experimental groups received a single trial of PAIRED presentation of AM and the sugar reward (AM+/EM or EM/AM+). The larvae in the control groups received neither the odour nor the sugar reward (HANDLING), only the SUGAR (EM+/EM or EM/EM+), or only the ODOUR (AM/EM or EM/AM). In the following test, the larvae were assayed for their preference for 1-OCT, determined with due adjustment according to Equation 1.

To quantify the impact of preconditioning as the difference between the SEPARATED and the COMPOUND cases, a difference index was calculated as:

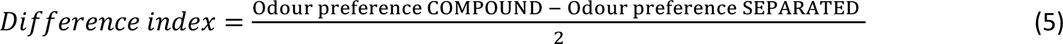

Accordingly, positive values indicate a higher odour preference in larvae that received the odours in COMPOUND during preconditioning as compared to larvae that received them SEPARATED; negative values indicate the opposite. Sensory preconditioning as an associative phenomenon would be indicated by higher difference indices in the experimental groups than in the controls.

For odour presentation, the Petri dish lids were equipped with four (during preconditioning and conditioning) or two (during the test) filter papers. One minute before the lid was placed on the Petri dish, the filter papers were loaded with 5 µl of AM and/or 5µl of 1-OCT. Other details were as described above. Variations on the paradigm are mentioned within the *Results* section.

#### Second-order conditioning

To test for second-order conditioning, the larvae first underwent a conditioning phase to establish an odour-reward association (first-order conditioning). This was followed by a second-order conditioning phase in which the previously rewarded odour was presented together with a novel, ‘target’ odour. Then the larvae were tested for their preference for the target odour (**Fig. 7**).

During first-order conditioning, the larvae in the experimental groups received a single PAIRED presentation of AM and the sugar reward (AM+/EM or EM/AM+). The larvae in the control groups received neither the odour nor the sugar reward (HANDLING), only the SUGAR (EM+/EM or EM/EM+), or only the ODOUR (AM/EM or EM/AM). This was followed by the second-order conditioning phase, during which the previously rewarded odour AM was presented either in COMPOUND with a novel, target odour (1-OCT) or temporally SEPARATED from the target odour; in both cases Petri dishes with agarose but no added sugar were used. To equate for the total duration of the experiment and for handling we added a blank trial for the COMPOUND case. In the following test, the larvae were assayed for their preference for 1-OCT, determined with due adjustment according to Equation 1.

To quantify the impact of the second-order conditioning phase, the difference between the SEPARATED and the COMPOUND cases was determined by calculating a difference index according to Equation 5. Positive values therefore indicate a higher odour preference in larvae that received the odours in COMPOUND during second-order conditioning as compared to larvae that received them SEPARATED; negative values indicate the opposite. Second-order conditioning as an associative phenomenon would be indicated by higher difference indices in the experimental groups than in the controls.

Other details were as described above. Variations on the paradigm are mentioned within the *Results* section.

### Statistics

Non-parametric statistics were performed throughout (Statistica 13, RRID:SCR_014213, StatSoft Inc., Tulsa, USA). To test whether values are significant relative to chance level (zero), one-sample sign tests (OSS) were used. To compare across multiple independent groups, Kruskal-Wallis tests (KW) with subsequent pair-wise comparisons by Mann-Whitney U-tests (MWU) were performed. To ensure a within-experiment error rate below 5%, Bonferroni-Holm corrections (Holm 1979) were applied. Data are shown as box plots with the median as the middle line, the 25 and 75 % quantiles as box boundaries, and the 10 and 90 % quantiles as whiskers.

The results of the statistical tests and the source data of all experiments are documented in the Data file **Supplemental_Table_S1.xlsx**.

## Results

### PAIRED and UNPAIRED training modulate odour preferences in an opposing manner

Odour preferences after one trial of PAIRED odour-reward training are higher than after UNPAIRED training, that is after presentations of odour temporally separated from the reward (**Fig. 1A**). Such a difference indicates appetitive associative memory and is reflected in a positive memory score (**Fig. 1B**). Corresponding results were obtained after three training trials (**Fig. 1C, D**), confirming earlier reports (for example Saumweber et al. 2011; El-Keredy et al. 2012; Schleyer et al. 2015a; Weiglein et al. 2019). However, such results do not allow one to conclude whether learning has taken place through PAIRED training, through UNPAIRED training, or both. This is because through PAIRED training the larvae may learn that the odour predicts the occurrence of the reward, leading to an increase in odour preference. Conversely, through UNPAIRED training they may learn the opposite, namely that the odour predicts the non-occurrence of the reward, leading to a decrease in odour preference. This begs the question as to what the baseline level of odour preference is, cleared of the associative effects of the training experience.

In larval *Drosophila* such a baseline odour preference can be conveniently determined by testing the animals in the presence of the reward. That is, the difference in odour preference after PAIRED versus UNPAIRED training that is observed under conventional testing conditions is abolished when the testing is carried out in the presence of the reward (**Fig. 2A**), leading to memory scores indistinguishable from chance level (**Fig. 2B**). This effect has been reported before, including controls showing that ‘innate’ odour preferences in experimentally naïve animals are not altered by the presence of the reward (for example Schleyer et al. 2011; Schleyer et al. 2015a; Paisios et al. 2017; Schleyer et al. 2020), a result that we replicate here (**Supplemental_Fig_S1.pdf**). These findings can be grasped best by the notion of appetitive associative memory supporting a learned search for the reward, which is adaptively abolished when the sought-for reward is indeed present (for more detailed discussions see Gerber and Hendel 2006; Schleyer et al. 2011). For the current context, the important point is that the residual odour preference that is observed when the reward is present during testing thus reflects the odour preference cleared of the influence of associative memories and can therefore be used as a baseline against which the associative effects of PAIRED and UNPAIRED training can be measured (stippled line in **Fig. 2A**). This shows that odour preferences after PAIRED training are increased relative to baseline, whereas after UNPAIRED training they are decreased (**Fig. 2A**). The same is observed after three training trials (**Fig. 2C, D**), confirming earlier results (Saumweber et al. 2011; Schleyer et al. 2011, 2015b; Paisios et al. 2017; Weiglein et al. 2019; Schleyer et al. 2020) (**Fig. 3**). We conclude that PAIRED versus UNPAIRED training have parametrically opposite effects on odour preference.

**Figure 2.**
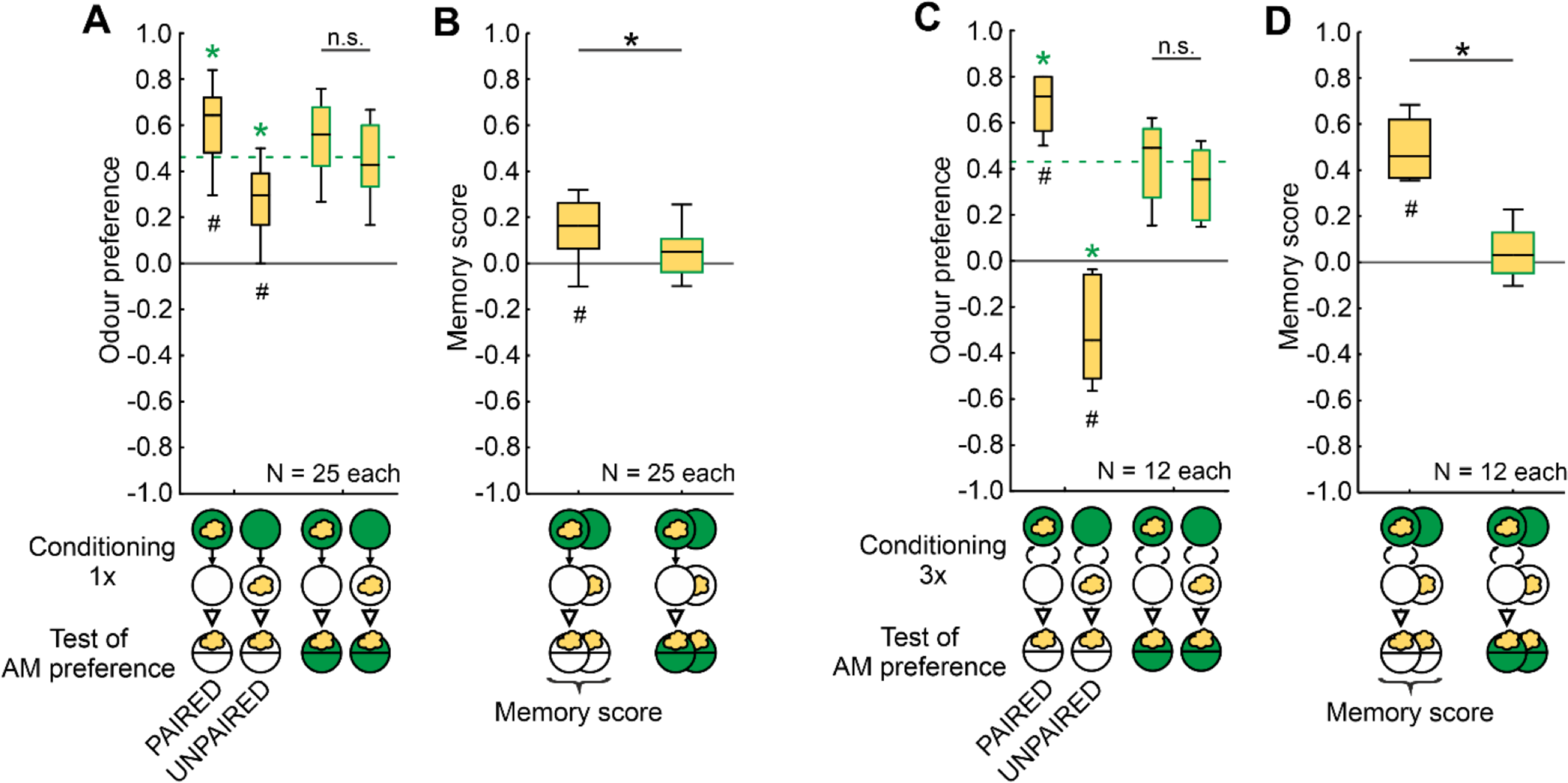
PAIRED and UNPAIRED training modulate odour preferences in an opposing manner. **(A-D)** Larvae were trained with PAIRED or UNPAIRED presentations of odour and the sugar reward and tested for their odour preference, either in the absence or in the presence of the reward. **(A)** After one-trial PAIRED training and when tested in the absence of the reward, the larvae show higher odour preferences than after UNPAIRED training (two box plots to the left). However, when the larvae were tested in the presence of the reward, this difference is abolished (two box plots to the right); their respective odour preference values can thus be pooled to represent baseline levels of odour preference, cleared of the influence of associative memories (green stippled line). Differences from this baseline demonstrate increased odour preferences after PAIRED and decreased odour preferences after UNPAIRED training. **(B)** Accordingly, memory scores are higher when the animals are tested in the absence than in the presence of the reward (innate odour preferences are unaffected by the presence of the reward: **Supplemental_Fig_S1.pdf**). **(C, D)** Corresponding results are observed after three training trials. Note the odour avoidance that is observed after UNPAIRED training when the test is carried out in the absence of the reward (C, second from left). Box plots represent the median, 25/75% and 10/90% quantiles. Sample sizes are indicated within the figure. # indicates significance from zero in an OSS test, * and n.s. indicate significance and non-significance, respectively, in an MWU-test. Specifically, a green * symbol refers to significance in an MWU-test against baseline, that is the pooled preferences when tested in the presence of the reward. Statistical results and source data in **Supplemental_Table_S1.xlsx**. Other details as in **Fig. 1**.

**Figure 3.**
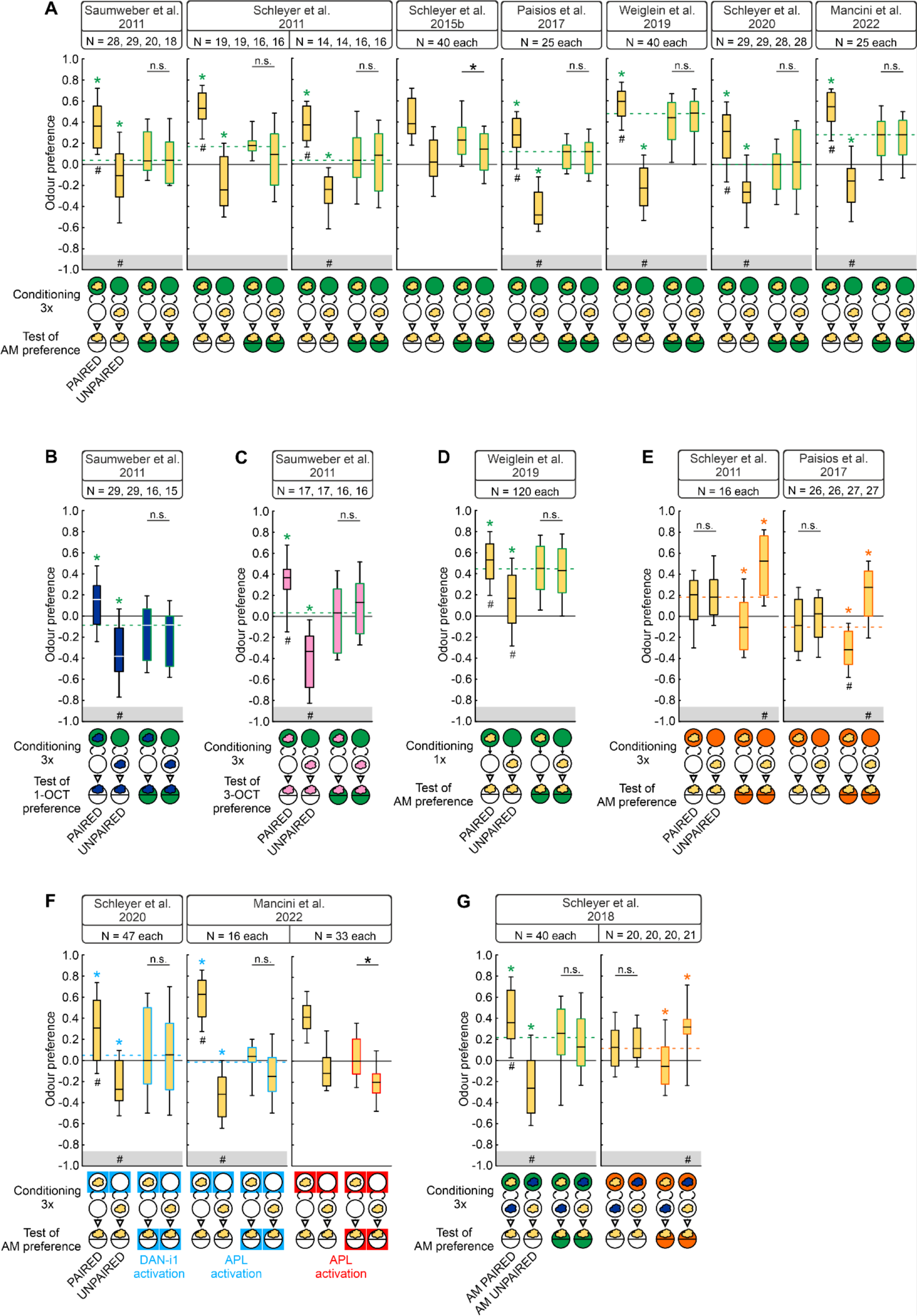
UNPAIRED training with reward can establish odour avoidance. **(A-G)** Survey of previously published experiments that revealed both PAIRED and UNPAIRED memory against baseline odour preferences (stippled lines) using different odours (clouds) (yellow, n-amylacetate; blue, 1-octanol; pink, 3-octanol) and rewards (circles/squares) (fructose, green; optogenetic activation of the indicated neurons, blue/red for ChR2-XXL/ Chrimson), or quinine as a punishment (orange). As indicated, training involved either one-odour ‘absolute’ conditioning (A-F), or two-odour differential conditioning followed by single-odour testing (G), with the indicated number of training trials. Of note is the variation in baseline odour preferences between experiments, arguably reflecting variation between for example seasons, food, experimental settings and experimenters, even when the same odour and concentration are used (also see **Supplemental_Fig_S2.pdf**). This demonstrates the importance of determining the baseline odour preferences separately for each experiment. Considering the total of 15 reward cases, in 11 out of the 13 datasets in which the baseline could be determined, UNPAIRED training established odour avoidance (85%). This argues for conditioned inhibition rather than learned inattention as the psychological mechanism for UNPAIRED memory. In two of the reward cases the baseline could not be determined because there were significant differences between the PAIRED and UNPAIRED trained groups tested in the presence of the reinforcer; this is indicated by the absence of the grey horizontal boxes at the bottom of the panels. For the special case of one-trial reward training see the discussion in the body text and **Fig. 4C**. The three cases of punishment reveal that UNPAIRED training established increased odour attraction, likewise arguing for conditioned inhibition rather than learned inattention. Critical evidence for conditioned inhibition is highlighted by placing the # symbol in the grey box at the bottom of the panels. Box plots represent the median, 25/75% and 10/90% quantiles. Sample sizes are indicated within the figure. # indicates significance from zero in an OSS test, * and n.s. indicate significance and non-significance, respectively, in an MWU-test. Coloured * symbols indicate significance in an MWU-test against baseline, that is against the pooled preferences when tested in the presence of the reinforcer. Statistical results and source data in **Supplemental_Table_S1.xlsx**. Other details as in **Fig. 1**.

**Figure 4.**
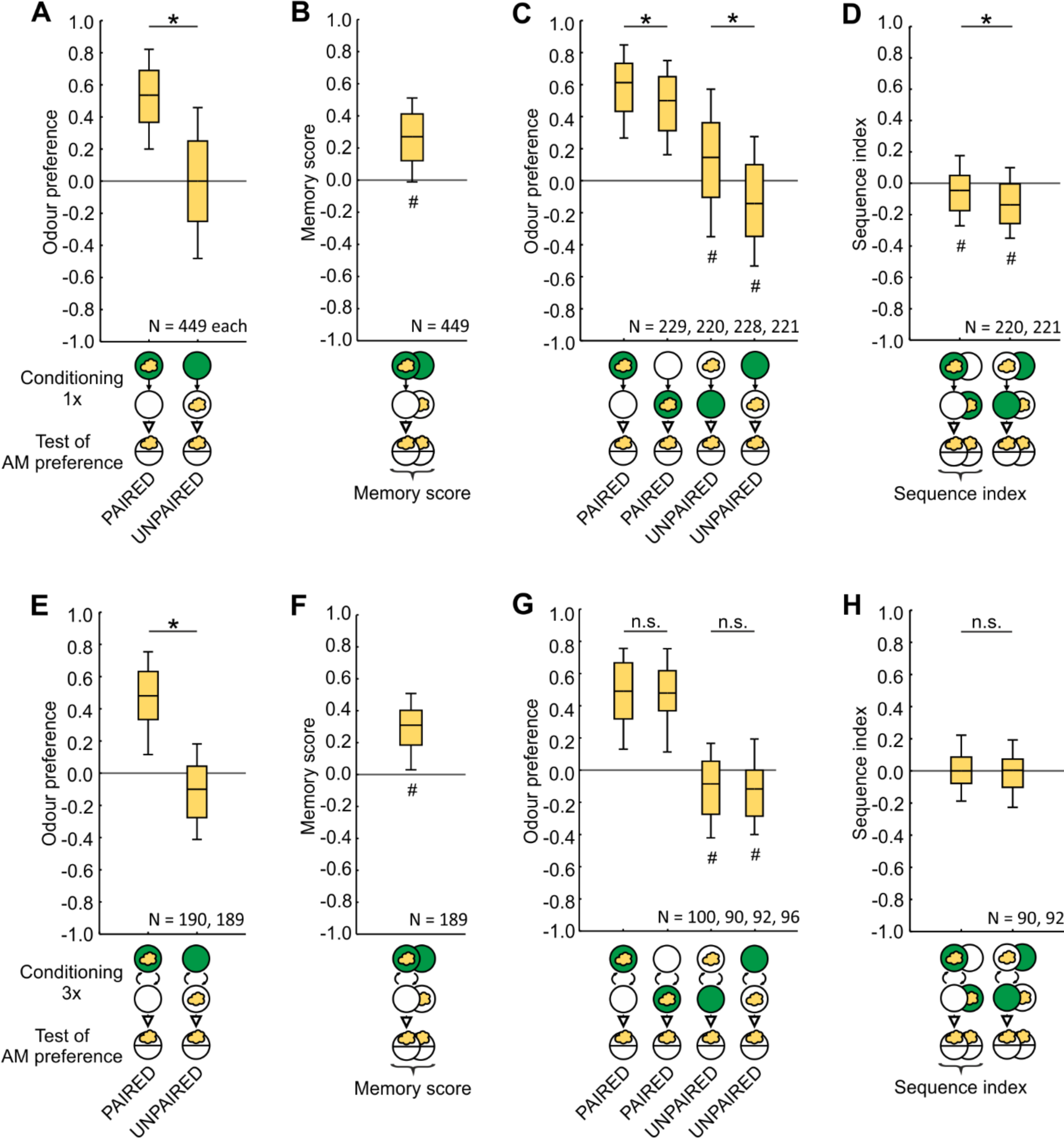
Stronger effect of the sequence of training events for UNPAIRED than for PAIRED training. **(A, B)** Larvae received one trial of either PAIRED or UNPAIRED training of odour and the sugar reward. The sequence of events during training was as depicted in half of the cases and the reverse in the other half of the cases (not shown here, but see C, D). In all cases, training was followed by a test of odour preference, **(A)** which turned out to be higher after PAIRED training than after UNPAIRED training, **(B)** resulting in positive memory scores indicative of appetitive associative memory. **(C, D)** Data from (A), separated by the sequence of events during training. **(C)** Upon PAIRED training, odour preference values are slightly lower when the blank period (open circle) comes first and the odour-reward pairing comes second, as compared to the reverse sequence (two box plots to the left). The same effect is observed upon UNPAIRED training (two box plots to the right). **(D)** A quantification of these differences by the sequence index (Equation 3, Equation 4) reveals stronger effects of the sequence of training events during UNPAIRED training as compared to PAIRED training. **(E-H)** As in (A-D), but for three training trials, **(E, F)** providing evidence for associative memory, **(G, H)** but no evidence of an effect of the sequence of events during training. Box plots represent the median, 25/75% and 10/90% quantiles. Sample sizes are indicated within the figure. # indicates significance to chance level, * and n.s. significance and non-significance, respectively, in an MWU-test. Statistical results are given along with source data in the data file **Supplemental_Table_S1.xlsx**. For data separated by individual dataset see **Supplemental_Fig_S2.pdf**. Other details as in **Fig. 1**.

**Figure 5.**
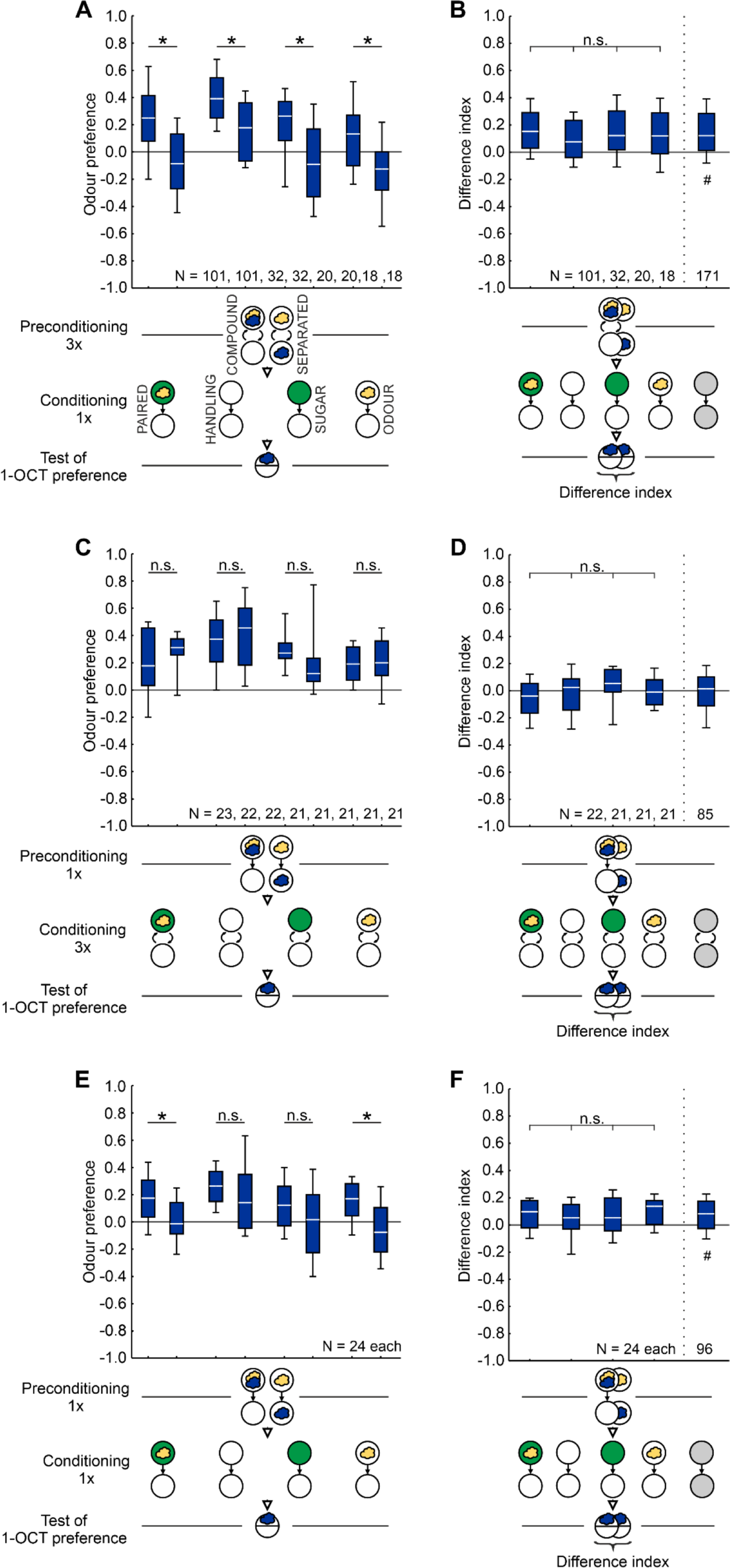
No evidence of sensory preconditioning. **(A-F)** During preconditioning the larvae were presented with two odours, AM (yellow cloud) and 1-OCT (blue cloud), either as a COMPOUND on the same Petri dish (white circle) or SEPARATED. During conditioning the larvae either received PAIRED training of the indicated odour and sugar reward (green circle), or they received associative controls during which either no odour and no sugar reward (HANDLING), only the SUGAR, or only the ODOUR were presented. Subsequently, the larvae were tested for their preference for the odour not used during the conditioning phase. Positive and negative preference scores reflect approach towards or avoidance of the odour in the test, respectively (Equation 1). The difference index quantifies the differences in odour preference between the respective COMPOUND and the SEPARATED group (Equation 5). Sensory preconditioning as an associative phenomenon would be indicated if the difference indices in the PAIRED group were higher than in the controls. No evidence for such a result was obtained in three variations on the experiment with the number of trials during the preconditioning/condition phases set to 3/1 **(A, B)**, 1/3 **(C, D)**, or 1/1 **(E, F)**. Box plots represent the median, 25/75% and 10/90% quantiles. Sample sizes are indicated within the figure; sample sizes for the PAIRED groups in (**A, B**) are high because, unlike all the other experiments in the present study, the control experiments were performed in a staggered, successive manner with the PAIRED groups always run in parallel. # indicates significance from zero in an OSS test, * and n.s. indicate significance and non-significance, respectively, in an MWU-test or KW-test. Statistical results and source data in **Supplemental_Table_S1.xlsx**. Other details as in **Fig. 1**.

### Does UNPAIRED training establish conditioned inhibition or learned inattention?

The opposing behavioural effects of PAIRED versus UNPAIRED training conform to the experimental psychology constructs of conditioned excitation versus conditioned inhibition and reflect the prediction of reward occurrence versus reward non-occurrence in prediction-error learning rules (for example Rescorla and Wagner 1972; Malaka 1999). However, an alternative scenario has been suggested whereby the respective training protocols result in increases versus decreases in how effectively the conditioned stimuli are processed (Mackintosh 1975; Pearce and Hall 1980). In the current case, this would suggest that the opposing effects of PAIRED versus UNPAIRED training are due to increased versus decreased learned attention to the odour. However, our results show that UNPAIRED training not only establishes decreases in odour preference relative to baseline, but indeed establishes avoidance (**Fig. 2C**). This confirms earlier results in larval *Drosophila* (**Fig. 3**) and is consistent with recent findings in adult flies (Jacob and Waddell 2020). Critically, these results are incompatible with learned inattention as an explanation for the effects of UNPAIRED training, because a lack of attention may reduce odour preference to zero, but cannot establish avoidance. This makes it superfluous to run summation tests as devised by Rescorla (1969) that check for learned inattention as an alternative scenario to explain learning through UNPAIRED training.

According to the prediction-error learning rules mentioned in the preceding paragraph, conditioned excitation ensues when there is a positive prediction error, that is when for example a reward is received although it was not predicted. In the current case, this is straightforward for PAIRED training when the odour is presented together with the reward. The prediction error is strongly positive in the first trial (‘pleasant surprise’) and becomes less and less as training progresses – because the reward is predicted better and better. This results in strong learning in the first trial and in an asymptotic learning curve as training progresses (Neuser et al. 2005; Weiglein et al. 2019).

Conversely, conditioned inhibition ensues when there is a negative prediction error, that is when for example a predicted reward is not actually received (‘frustrating surprise’). For UNPAIRED training, this is not straightforward, however. In the current case, this is because one wonders what the source of a reward-prediction would be at the moment of odour presentation. Prediction-error learning rules typically make the assumption that presenting the reward alone can establish associative memories for the context (Dweck and Wagner 1970; Rescorla 1972; Bouton and Nelson 1998). When the odour is subsequently presented in that same context, these context-reward associations provide an expectation of reward which, however, is frustratingly not received. The ensuing negative prediction error is then the basis for conditioned inhibition to accrue to the odour. This provides an interesting experimental test because accordingly negative prediction errors would only arise when during UNPAIRED training the reward-only presentation comes first, establishing context-reward associations, and the odour-only presentation comes second. For the reverse sequence of the odour being presented first, there would not yet be any context-reward association that could be frustrated. Such sequence dependence of odour preference values after UNPAIRED training should be particularly prominent when only one training trial is used, because for multiple training trials the contextual memories established in trial #1 would need to be reckoned with from trial #2 on, effectively ‘ironing out’ sequence-effects as training proceeds. Notably, the learned inattention scenario proposed to account for learning through UNPAIRED training would not suggest sequence-effects in either case. With these considerations in mind, we analysed two large datasets that we have accumulated over the years with course students, interns, scientific guests and apprentice staff using our standard one-trial training procedure (**Fig. 4A-D**; total sample size N = 898) as well as corresponding experiments with three training trials (**Fig. 4E-H**; total sample size N = 379).

These combined datasets confirm that odour preference values after one-trial PAIRED training are higher than after UNPAIRED training (**Fig. 4A**), yielding a positive memory score indicative of appetitive associative memory (**Fig. 4B**); corresponding results were obtained for the dataset using three training trials (**Fig. 4E, F**). When odour preference values are separated by the sequence of odour and sugar reward presentation during training, it turns out that for the PAIRED case of one-trial training, odour preference values are slightly and, given the large sample size, significantly lower when the blank presentation (EM) comes first and the odour-reward pairing (AM+) comes second as compared to the reverse AM+/EM sequence (**Fig. 4C, leftmost two box plots**). The evidence is more clear-cut for the UNPAIRED case, with odour preference values not only lower for the EM+/AM sequence than for the AM/EM+ sequence (**Fig. 4C, rightmost two box plots**), but critically showing odour avoidance for only the EM+/AM sequence. Compared to the PAIRED case, the sequence of training events has more impact in the UNPAIRED case, as shown by differences in the sequence index (**Fig. 4D**). No differences in odour preference values between the training sequences were observed for three-trial training (**Fig. 4G**, **H**).

These results conform to prediction-error learning rules but cannot be explained by recourse to changes in learned attention as the psychological mechanism for UNPAIRED learning.

### No evidence of sensory preconditioning

Larvae first received two odours (AM and 1-OCT) either as a COMPOUND or SEPARATED from each other (preconditioning phase, 3 trials). For both groups this was followed by PAIRED odour-sugar reward training with one of the odours (AM+) (conditioning phase, one trial) and testing with the other odour (1-OCT) (**Fig. 5A**). We reasoned that if there is sensory preconditioning then the larvae of the COMPOUND group should show higher odour preference than the larvae of the SEPARATED group, as was indeed the case (**Fig. 5A**).

However, the same difference in odour preference is observed in control groups for which the treatment in the conditioning phase was varied such that no odour-sugar reward association could be formed (**Fig. 5A**). Importantly, the difference indices did not vary across the experimental and control groups (**Fig. 5B**). Thus, although experiencing the two odours in COMPOUND during preconditioning resulted in consistently higher odour preference during the test than experiencing them SEPARATED from each other, these results do not provide evidence of sensory preconditioning as an associative phenomenon.

Reducing the number of trials during preconditioning to one and increasing the number of trials during conditioning to three did not provide evidence for sensory preconditioning, either. In this case, the groups that received the odours in COMPOUND during preconditioning and those that received them SEPARATED showed statistically indistinguishable odour preferences regardless of the treatment during conditioning (**Fig. 5C**), resulting in equal difference indices (**Fig. 5D**). The difference indices were likewise indistinguishable across experimental and control groups when only one trial was used for both preconditioning and conditioning (**Fig. 5E, F**).

Given that, surprisingly as we hesitate to add, the treatment during conditioning did not make any difference in the above experiments, we wondered whether under the current regimen an association between the odour (AM) and the sugar reward was indeed established during conditioning, and whether it was indeed behaviourally effective during the test. The larvae first underwent three trials of preconditioning with the two odours presented in COMPOUND or SEPARATED, followed by a single conditioning trial with PAIRED or UNPAIRED training of the odour AM and the sugar reward. We then tested the animals for their AM preference and found these to be higher after PAIRED than after UNPAIRED training, indicating effective association formation (**Fig. 6A**) and resulting in positive memory scores (**Fig. 6B**). Interestingly, the memory scores were not different between the animals that had received the two odours in COMPOUND or SEPARATED during preconditioning, meaning that the type of odour exposure during preconditioning did not affect association formation during the conditioning phase.

**Figure 6.**
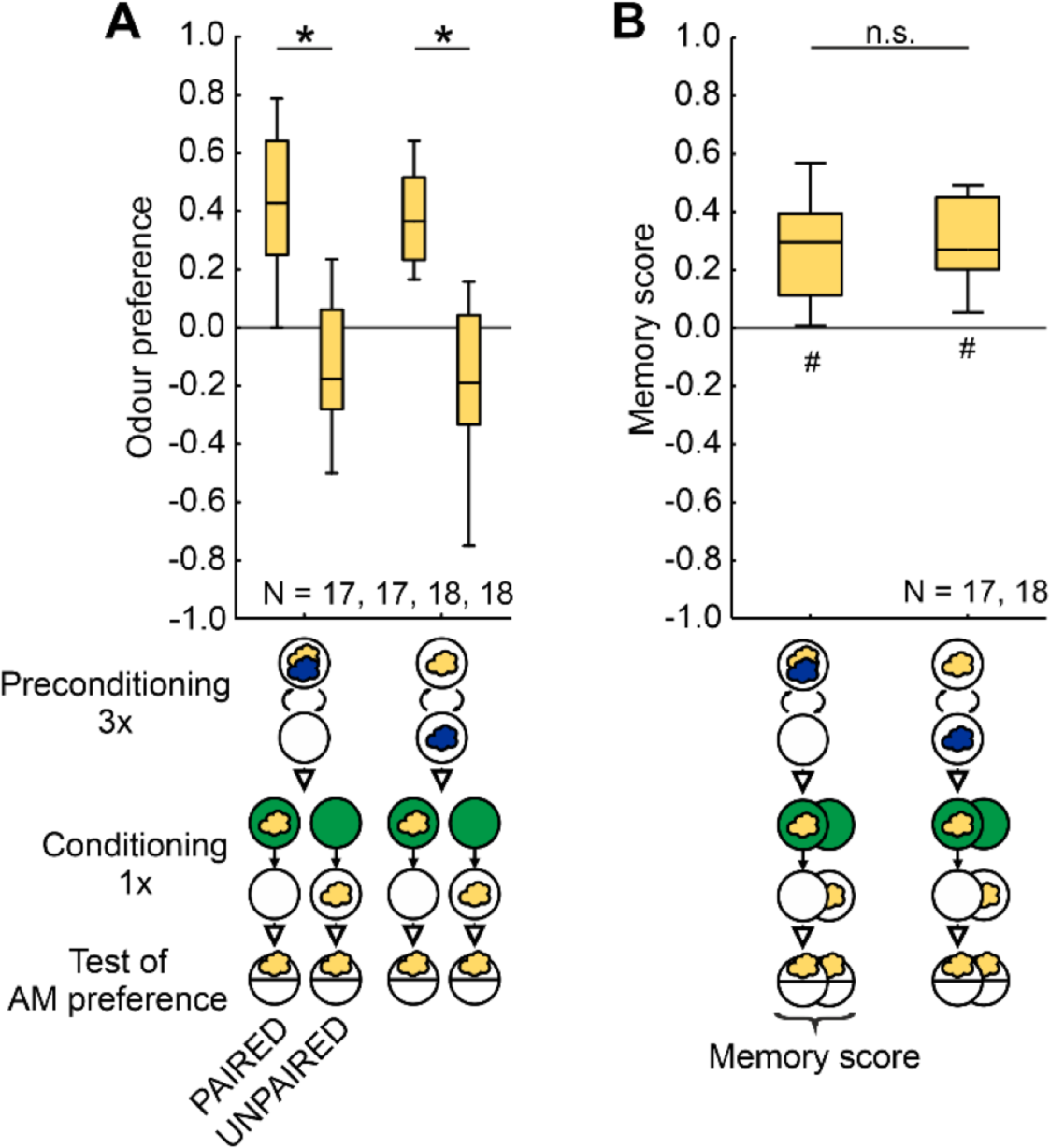
Effective association formation during the conditioning phase. **(A-B)** During preconditioning the larvae received the two odours either in COMPOUND or SEPARATED. This was followed by either PAIRED or UNPAIRED conditioning of the indicated odour and the sugar reward. Subsequently the larvae were tested for their preference for the odour used during conditioning. **(A)** Regardless of the type of preconditioning, PAIRED training led to higher odour preferences than UNPAIRED training, indicating effective association formation during conditioning. **(B)** Memory scores did not differ between groups, indicating that association formation during conditioning was equally effective regardless of the preconditioning treatment. Box plots represent the median, 25/75% and 10/90% quantiles. Sample sizes are indicated within the figure. # indicates significance from zero in an OSS test, * and n.s. indicate significance and non-significance, respectively, in an MWU-test. Statistical results and source data in **Supplemental_Table_S1.xlsx**. All other details as in **Fig. 5**.

**Figure 7.**
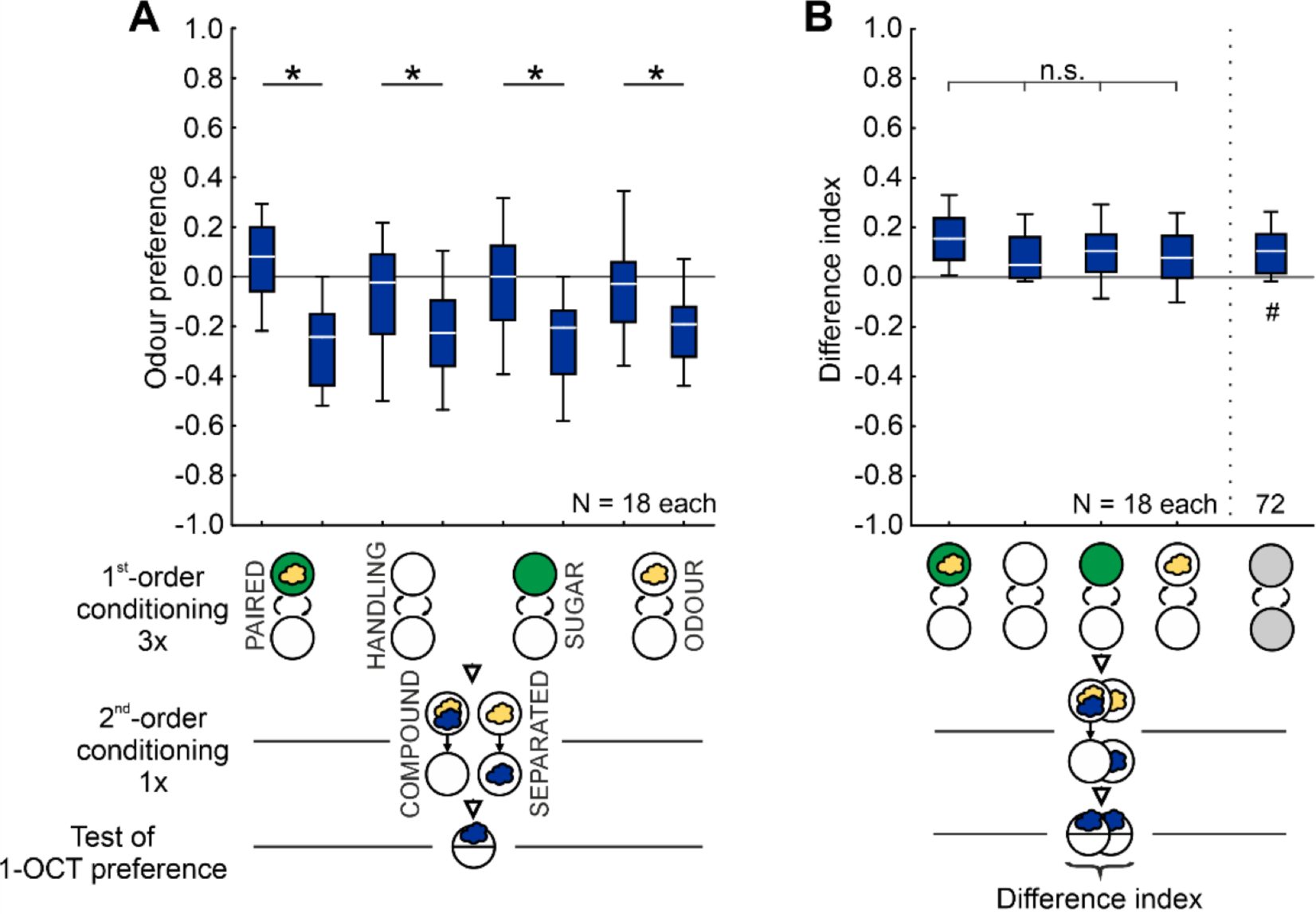
No evidence of second-order conditioning. **(A-B)** During first-order conditioning, the larvae in the experimental groups received three trials of PAIRED training of AM (yellow cloud) and a sugar reward, whereas control groups received either no odour and no sugar reward (HANDLING), only the SUGAR, or only the ODOUR. In the following, second-order conditioning phase, the larvae were presented with AM either in COMPOUND with or SEPARATED from a novel, ‘target’ odour (1-OCT) (blue cloud). Subsequently, the larvae were tested for their preference for the target odour. Second-order conditioning as an associative phenomenon would be indicated if the difference indices, quantifying the effect of the second-order conditioning phase, were higher in the experimental than in the control groups. No evidence for such a result was obtained. Box plots represent the median, 25/75% and 10/90% quantiles. Sample sizes are indicated within the figure. # indicates significance to chance level, * and n.s. significance and non-significance, respectively, in MWU-tests or a KW-test. Statistical results and source data in **Supplemental_Table_S1.xlsx**. All other details as in **Fig. 5**.

Taken together, these results do not offer evidence of sensory preconditioning as an associative phenomenon in larval *Drosophila*.

### No evidence of second-order conditioning

Larvae in the experimental groups first underwent conditioning to establish an odour-reward association (first-order conditioning, 3 trials). This was followed by a second-order conditioning phase in which the previously rewarded odour was presented either in COMPOUND with or SEPARATED from a novel, target odour (1 trial). Then the larvae were tested for their preference for the target odour (**Fig. 7**). We reasoned that if there is second-order conditioning then the larvae of the COMPOUND group should show a higher odour preference than the larvae of the SEPARATED group, as was indeed the case (**Fig. 7A**). However, the same difference in odour preference is observed in control groups for which the treatment during first-order conditioning was varied such that no odour-sugar reward association could be formed (**Fig. 7A**). Importantly, the difference indices did not vary across the experimental and control groups (**Fig. 7B**). These results do not provide evidence of second-order conditioning as an associative phenomenon (preliminary experiments in the aversive domain, using a modified protocol and associations between odour and electric shock during first-order conditioning, likewise yielded no such evidence: **Supplemental_Fig_S3.pdf**).

To test for the possibility that the treatments in the second phase of the experiment rendered the first-order association ineffective, we conducted a control experiment in which the larvae were trained either PAIRED or UNPAIRED with the first-order conditioning odour (AM) and a sugar reward, followed by presentation of the first-order conditioning odour either in COMPOUND with or SEPARATED from the target odour (1-OCT). This was followed by testing for the preference for the first-order conditioning odour (**Fig. 8A**). PAIRED training with the first-order conditioning odour yielded higher preference for it than UNPAIRED training with it, both when it was subsequently presented in COMPOUND with and SEPARATED from the target odour (**Fig. 8A**). Accordingly, appetitive associative memory was observed in both cases (**Fig. 8B**), showing that the effects of first-order conditioning were not rendered ineffective by the treatments in the second experimental phase. We note that, when compared to the COMPOUND case, the memory scores were less when the two odours were presented SEPARATED in the second-order conditioning phase, arguably because extinction learning is more effective when the first-order conditioning odour is experienced alone during this part of the experiment. In any event, the conclusion remains that from the present results there is no evidence of second-order conditioning as an associative phenomenon in larval *Drosophila*.

**Figure 8.**
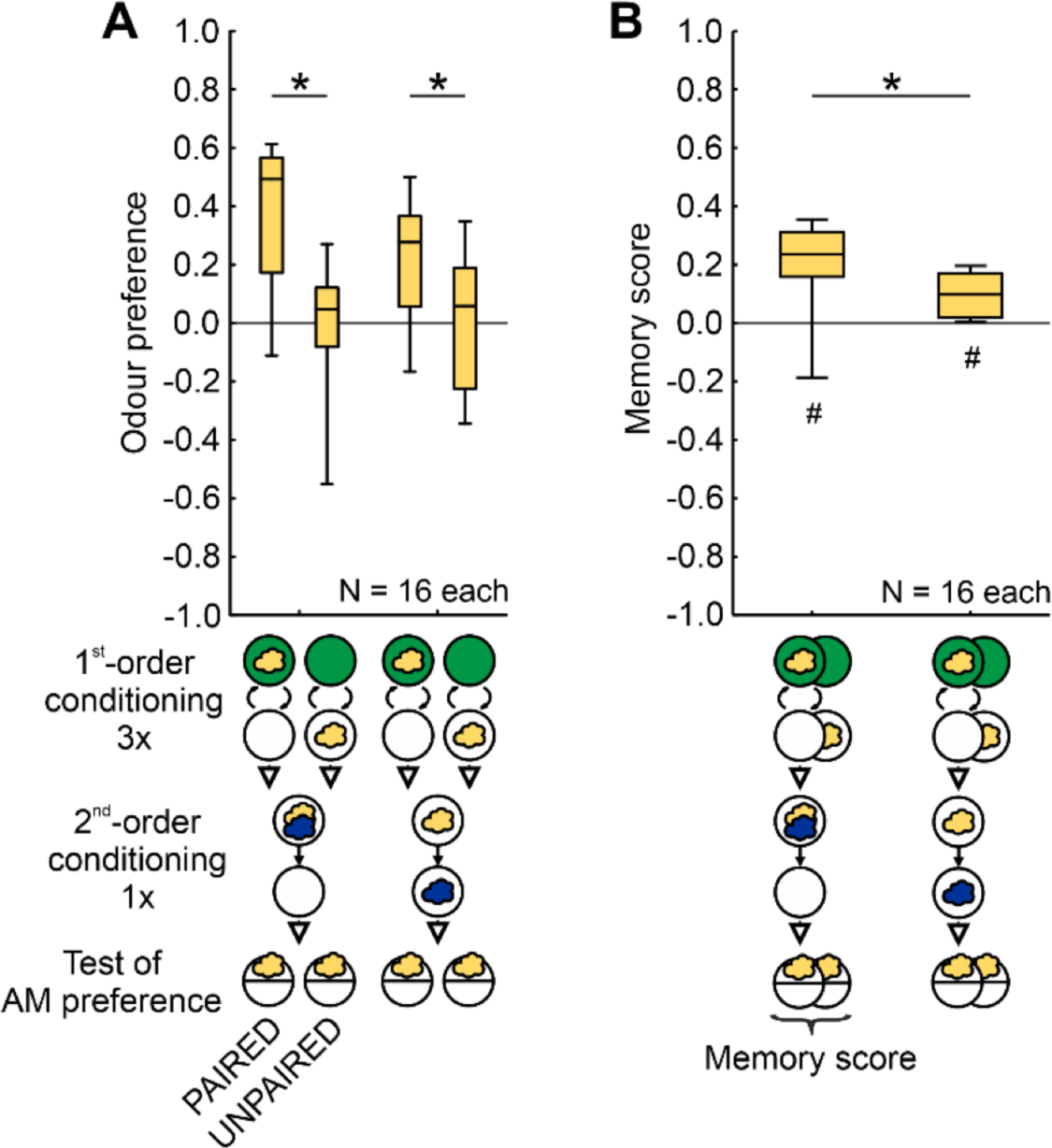
Effective first-order associations. **(A-B)** Larvae received three trials of first-order conditioning with either PAIRED or UNPAIRED training of AM (yellow cloud) and a sugar reward. This was followed by presenting AM either in COMPOUND with or SEPARATED from 1-OCT (blue cloud) as the target odour, followed by testing for the preference for AM as the first-order conditioning odour. Regardless of the type of odour presentation in the second experimental phase, PAIRED training led to higher odour preferences than UNPAIRED training, resulting in positive memory scores in both cases and indicating effective first-order conditioning associations. In comparison to the COMPOUND case, the memory scores were less when the two odours were presented SEPARATED, though, arguing for more effective extinction learning when the first-order conditioning odour is experienced alone. Box plots represent the median, 25/75% and 10/90% quantiles. Sample sizes are indicated within the figure. # indicates significance to chance level, * significance in an MWU-test. Statistical results and source data in **Supplemental_Table_S1.xlsx**. All other details as in **Fig. 5**.

## Discussion

The present data provide evidence in support of conditioned inhibition in larval *Drosophila*, corresponding to what has been reported in adult flies (Barth et al. 2014; Jacob and Waddell 2020) and honeybees (Takeda 1961; Bitterman et al. 1983; Chandra et al. 2010; Matsumoto et al. 2012). Accordingly, many models of learning account for conditioned inhibition (see section *Maggots versus models*). However, our data do not provide evidence for sensory preconditioning or second-order conditioning in larval *Drosophila*. Such absence of evidence does not amount to evidence of absence, though. Changes to the experimental protocol that could uncover these forms of learning in larval *Drosophila* include changes to the number or temporal spacing of trials, the use of different odour concentrations, different odours or combinations of odours with visual stimuli, or other kinds of reinforcer. Also, more fine-grained analyses of behaviour through video tracking might reveal effects of training that the current study has overlooked. It is with these caveats in mind that the following discussion supposes that larval *Drosophila* are not capable of sensory preconditioning and second-order conditioning. Why not?

### Maggots versus adult flies and honeybees: Ways of life

Sensory preconditioning and second-order conditioning have been demonstrated in adult flies (Brembs and Heisenberg 2001; Guo and Guo 2005; Tabone and De Belle 2011; Martinez-Cervantes et al. 2022; Yamada et al. 2023) and honeybees (Takeda 1961; Bitterman et al. 1983; Müller et al. 2000; Hussaini et al. 2007), including with protocols that use binary odour compounds. In contrast, we report an absence of evidence for these forms of learning in larval *Drosophila*. Which of the differences in the biology of larval *Drosophila* versus adult flies and honeybees could account for this discrepancy?

In insects with complete metamorphosis the larval stage is sexually undifferentiated and is typically dedicated to feeding and growth. Indeed, the salivary glands of larval *Drosophila* are larger than their brain. Foraging worker honeybees, the caste of bee used in the above-cited experiments, are likewise asexual in motivation and, as their designation betrays, dedicated to foraging, too. In motivational terms, therefore, larval *Drosophila* seem closer to honeybees than to adult flies.

What differentiates larval *Drosophila* from adult flies and honeybees is rather that the larvae live on or in decaying fruit as their ‘food-home’. They thus do not need to bother much with finding as-yet absent resources such as food, a home (as is the case for honeybees), or mating partners (as adult flies need to do). It would seem that larvae can therefore organize their behaviour in a more ‘online’ manner, that is through sensory-motor loops in relation to physically present resources. For adult flies and honeybees, in contrast, behaviour needs to be organized in a more ‘offline’ manner, towards desired but as-yet physically absent goals. As argued in the *Introduction* section, sensory preconditioning and second-order conditioning both require chained processing in relation to physically absent cues or reinforcement and thus chained ‘offline’ processing. These tasks might therefore tap into cognitive abilities bound up with the way of life of adult flies and honeybees. Does the organization of the brains of larval *Drosophila* versus adult flies and honeybees reflect this situation?

### Maggots versus adult flies and honeybees: Brain organization

The brains of larval *Drosophila*, adult flies, honeybees and indeed insects more generally are organized in largely the same way, including the mushroom body, the central-brain structure serving odour-tastant associative memory formation (for a strongly simplified overview see **Fig. 9A, B**). It therefore seems that the relevant differences in brain organization between larval *Drosophila* versus adult flies and honeybees are not to be found in the principles but in the details (Menzel 2013). Which details and in which brain structures?

**Figure 9.**
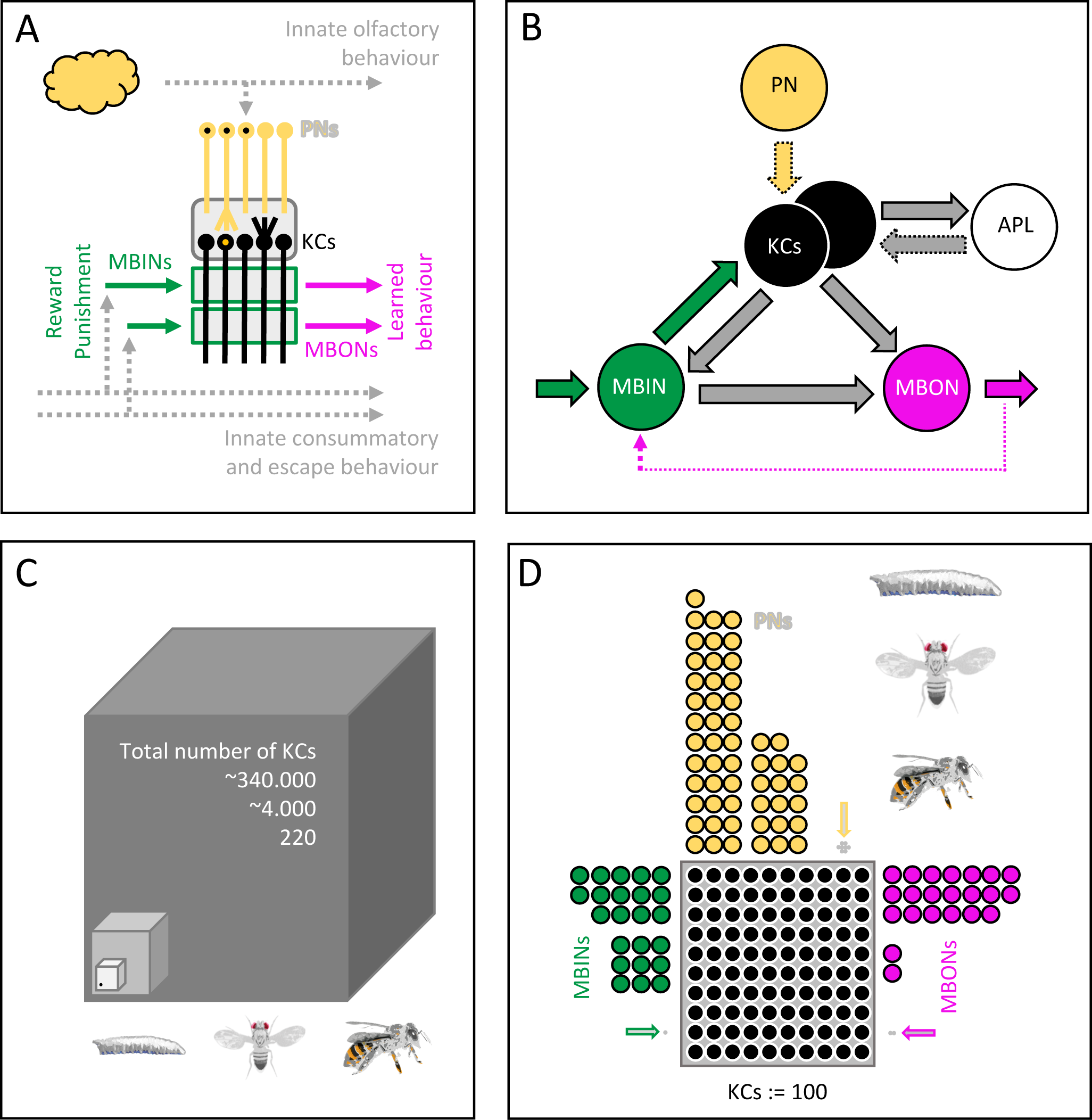
Simplified organization of the mushroom body. **(A)** The mushroom body intrinsic neurons (Kenyon cells, KCs; black) form a sparse, combinatorial representation of odours, established through a divergence/ convergence connectivity with sensory projection neurons (PNs; yellow) in the calyx (rounded grey rectangle). Axonal projections of the KCs pass through compartments (green rectangles) established by axonal branches of modulatory mushroom body input neurons (MBINs; green) (most of these are dopaminergic) and dendritic branches of mushroom body output neurons (MBONs; pink). Of a total of ten compartments, one with a rewarding and one with a punishing MBIN is shown. Associative memories are established through compartmentally specific plasticity at the synapses between odour-activated KCs and MBONs. This changes the balance of activity in avoidance-versus approach-promoting MBONs underlying learned behaviour. Pathways for innate behaviour are shown in grey. **(B)** PN→KC synapses are found only in the input region of the mushroom body (the calyx), but not in the lobe compartments (stippled yellow arrow). In larval *Drosophila*, APL→KC synapses are found only in the calyx (stippled grey arrow), rather than throughout the mushroom body as in adult flies; also, only six out of ten compartments, rather than all compartments as in adult flies, feature KC→APL synapses. The stippled pink arrow indicates indirect feedback from MBONs to MBINs. Innervations by more than one MBIN or MBON, multiple-compartment MBINs and MBONs, and lateral connections between MBONs as well as between KCs are not shown. **(C-D)** Cell numbers in the mushroom bodies of larval *Drosophila*, adult flies and honeybees summed for both hemispheres. Shown in (C) are the numbers of KCs (the dot indicates a single KC within a cube of 6×6×6 KCs), and in (D) the number of the indicated cell types normalized to the number of KCs. Numbers can be found in **Supplemental_Table_S1.xlsx** and are based on Eichler et al. (2017) and Saumweber et al. (2018) for larval *Drosophila*, and on Bates et al. (2020), Li et al. (2020) and Schlegel et al. (2021) for adult flies. Regarding honeybees, our estimations are based on Witthöft (1967), Schäfer and Rehder (1989), Maronde (1991), Rybak and Menzel (1993), Kreissl et al. (1994), Rybak (1994), Stevenson and Spörhase-Eichmann (1995), Grünewald (1999), Gronenberg (2001), Schröter and Menzel (2003), Schröter et al. (2007), Rybak (2011) as well as Zwaka et al. (2018).

CENTRAL COMPLEX. Although the brain of larval *Drosophila* is organized according to the same principles as in adult flies and honeybees, the numbers of neurons are reduced. Indeed, the central complex comprises so few neurons in the case of larval *Drosophila* that it cannot be recognized as a histological structure but only from the circuit motifs that these neurons establish. In contrast, the central complex of larger adult insects can be seen with the naked eye upon opening the head capsule and removing the surrounding tissue. Central complex function underlies course control relative to local sensory cues during the pursuit of as-yet absent goals (Pfeiffer 2023), for example when honeybees navigate between their hive and flower patches, sometimes covering kilometres and thus going way beyond the catchment area of their sensory systems. To the extent that a more sophisticated organization of behaviour in relation to absent goals is required for sensory preconditioning and second-order conditioning, the performance of these tasks might therefore come more easily to adult flies and honeybees with their more elaborate central complex.

ASCENDING OLFACTORY PATHWAYS. In comparison to adult flies and honeybees, the ascending olfactory pathways in larval *Drosophila* are characterized by lower numbers of expressed olfactory receptor genes, sensory neurons and projection neurons (PNs) and a lack of cellular redundancy at these layers of processing, as well as by less interhemispheric communication, a lower number of mushroom body intrinsic neurons (called Kenyon cells: KCs), and by the fact that about a fifth of these KCs receive input from only a single PN as part of what appears to be a labelled-line sensory-motor pathway. Together, this suggests that the ascending olfactory system has a substantially poorer signal-to-noise ratio and provides a much less nuanced representation of odours in larval *Drosophila* than in adult flies and honeybees. The present protocols for sensory preconditioning and second-order conditioning might thus be too demanding, as they require an ability to process binary odour compounds by their elements (something of which larvae are in principle capable, though: Chen and Gerber 2014; Chen et al. 2017) and a process of pattern completion when only one of these elements is physically present.

MUSHROOM BODY. The number of KCs is relatively similar in larval *Drosophila* and adult flies, at least in comparison to the much higher number in honeybees (**Fig. 9C**). Analysis of the number of PNs and the number of modulatory mushroom body input neurons (MBINs) *relative to the number of KCs* paints a similar picture (**Fig. 9D**). By the same token, the relative number of mushroom body output neurons (MBONs), respectively differing 10-fold between larval *Drosophila* and adult flies, as well as between adult flies and honeybees, does not account for why larval *Drosophila* should differ from both adult flies and honeybees in terms of sensory preconditioning or second-order conditioning.

APL and DPM. The mushroom body of *Drosophila* features a giant intrinsic neuron called APL. This GABAergic neuron is embryonically born, persists through metamorphosis, and functionally corresponds to the GABAergic A3 neurons in honeybees (Rybak and Menzel 1993). It is central to mushroom body function because it receives input from all KCs and provides inhibitory output back to them (Mancini et al. 2022 and references therein). In adult flies such reciprocal connectivity is observed throughout the mushroom body (as is the case for the KCs and GABAergic neurons in honeybees: Ganeshina and Menzel 2001; Zwaka et al. 2018). In larval *Drosophila*, however, reciprocal APL←→KC connectivity is only found in the calyx region of the mushroom body, where the signals from the PNs are received. In the lobe regions of the larval mushroom body, where all but two of the MBONs are located and where behavioural output is instructed, APL→KC connections are conspicuously absent. It would thus seem that in adult flies APL (and in the case of honeybees the A3LC neurons: Zwaka et al. 2018) can ‘take KCs offline’ from behavioural control whereas this would not be possible in larval *Drosophila* (for the significance of such offline processing for action planning and cognition see Menzel 2013). Interestingly, lobe-to-calyx feedback, possible through APL in larval *Drosophila* and adult flies and through the A3FB neurons in honeybees (Zwaka et al. 2018), was suggested to underlie reversal learning (Boitard et al. 2015) – a faculty indeed observed in all three kinds of animal (Tully and Quinn 1985; Boitard et al. 2015; Mancini et al. 2019). In addition, in adult flies APL is electrically coupled to the DPM neuron, which is absent in larval *Drosophila*, and concerted APL-DPM action has been reported to moderate KC-KC communication (Okray et al. 2022) in a process that should support pattern completion (Cayco-Gajic and Silver 2019). To the extent that mushroom body function ‘offline’ from behavioural control or pattern completion is involved in sensory preconditioning and second-order conditioning, the absence of these forms of learning in larval *Drosophila* could thus be related to differences in APL organization or the absence of DPM.

There thus seem to be four aspects of brain organization in larval *Drosophila* versus adult flies and honeybees that have the potential to explain the absence of sensory preconditioning and second-order conditioning in larvae: i) an insufficiently differentiated central complex; ii) an insufficient signal-to-noise ratio in the ascending olfactory pathways and thus an insufficiently nuanced representation of odour mixtures; iii) the absence of APL→KC connections in the mushroom body output region; and iv) the absence of the DPM neuron.

In the light of these considerations, we would like to note the complexities of behaviour that have nevertheless been observed in larval *Drosophila*. Beyond various forms of taxis and their state-dependent modulation (Vogt et al. 2021), these include associative learning between odours or visual cues and various kinds of taste reinforcement (Thum and Gerber 2019 and references therein), as well as discrimination, generalization, reversal learning, memory consolidation, and an adaptive dominance of consummatory over learned behaviour (Gerber and Hendel 2006; Mishra et al. 2010; Schleyer et al. 2011, 2015a; Widmann et al. 2016; Mancini et al. 2019).

### Maggots versus models

Conditioned inhibition through unpaired training of a cue with a reward can be accommodated by prediction-error learning rules (Malaka 1999 and references therein) (for a model inspired by the functional anatomy of the mushroom body not involving prediction errors: Gkanias et al. 2022). To account for conditioned inhibition, prediction-error models require that at the moment of presenting the cue there is a negative prediction error, as is the case when a predicted reward, frustratingly, is not actually received. This raises the question of the source of the reward prediction. Two non-exclusive scenarios have been proposed. A reward prediction may arise from previously established context-reward associations (Rescorla and Wagner 1972) or from the generalization of previously established cue-reward associations (Jürgensen et al. 2022). Our observation that unpaired training does not lead to odour avoidance when there is no possibility for context-reward associations to have been established (**Fig. 4C**, second box plot from the right) supports the contextual scenario.

Sensory preconditioning, however, is beyond the scope of the aforementioned models because it takes place between two cues rather than between a cue and reinforcement. What is required for sensory preconditioning is an experience-dependent process of configuration-learning, that is a process that endows two cues, based on their past co-occurrence, with the capacity to call up each otheŕs representation when presented alone. In analogy to pattern completion in the cerebellum and the hippocampus (Rudy and Sutherland 1995; Rolls 2013; Cayco-Gajic and Silver 2019), this could be achieved by modulations of KC-KC signalling (Müller et al. 2000; Okray et al. 2022). It would be interesting to see whether or not models of the mushroom body of larval *Drosophila* or adult flies that either do or do not feature the possibility of modulations in KC-KC signalling yield sensory preconditioning.

To account for second-order conditioning was one of the main goals in the development of real-time models of reinforcement learning through prediction errors (Malaka 1999 and references therein) (for an alternative scenario not involving prediction errors: Gkanias et al. 2022). What these prediction-error models have in common is that by virtue of the A+ association established in phase (i) cue A can call up a representation of +. During AB presentations in phase (ii) this associatively-activated representation of + can be associated with B. The resulting B+ association is thus the immediate cause of the response to B during the test. Given that a subset of dopaminergic mushroom body input neurons (MBINs of the DAN type) is thought to mediate internal reward signals, this raises the question of how a learned cue A can activate DANs. As associative learning entails plasticity at the KC→MBON synapses, this could happen through these same modulated synapses and feedback from the MBONs to the DANs. Such feedback was recently shown by Yamada et al. (2023) to underlie second-order conditioning in adult flies (see also König et al. 2019) and is compatible with the functional anatomy of the mushroom body in larval *Drosophila* as well (Eschbach et al. 2020). But why, then, is second-order conditioning not experimentally observed in the case of the larva?

Modelling studies typically restrict themselves to the task-relevant pathways. For example, they consider the ascending olfactory and gustatory pathways, the mushroom body, and the first steps of mushroom body efferent circuits. However, what goes on in the ‘rest’ of the animal is typically ignored, for example in relation to the visual, thermo-, mechano- or hygrosensory pathways, in relation to processing in the central complex or the ventral nerve cord, or in relation to signalling between brain, gut and glands (for an exception see Sakagiannis et al. 2021). The low number of neurons and the absence of cellular redundancy in larval *Drosophila* possibly makes them more susceptible to such influences than adult flies or honeybees. It would therefore be interesting to challenge models of the mushroom body that use numbers of model neurons corresponding to the case of larval *Drosophila* versus adult flies and honeybees with various levels of noise to see how this affects first- and second-order conditioning.

We note that there is a fundamentally different view of second-order conditioning, namely that it is a phenomenon of memory retrieval (Savastano and Miller 1998; Molet et al. 2012). It is suggested that during phase (ii) an AB association is formed and that through this AB association, *at the moment of the test*, B calls up A, which then calls up +. In other words, the immediate cause of the response is the A+ association established in phase (i), not a B+ association established in phase (ii). Such a retrieval-account of second-order conditioning has, to the best of our knowledge, not so far been related to the functional anatomy of the insect brain.

In closing, we would like to argue that when it comes to identifying the cognitive limits of larval *Drosophila* the apparent absence of sensory preconditioning and second-order conditioning is a success – because limits can only be determined through a failure to transgress them.

## Acknowledgements

Discussions with past and present members of our department and in particular with Michael Schleyer (U Sapporo, Japan) and Christian König (LIN) as well as Markus Fendt (U Magdeburg), technical assistance by Anna Ciuraszkiewicz-Wojciech, Bettina Kracht, and Thomas Niewalda (LIN), and support from the workshop of the Schwann-Schleiden Research Centre of the University of Göttingen are gratefully acknowledged. In addition to the course students, interns, scientific guests and apprentice staff that kindly worked with us on the experiments shown in **Figure 4**, we want to thank Fateme Asadzadeh, Arman Behrad, Niklas Behrenbruch, Finn Both, Vivian Brunsberg, Melissa Comstock, Luke Flanagan, Laura Kilian, Anna Matkovskaia, Celine Seiffert, and Esmeralda Tafani for contributing data, and Rupert Glasgow (Zaragoza, Spain) for English language advice. Special thanks go to Yoshinori Aso (HHMI Janelia), Randolf Menzel (FU Berlin), and Jürgen Rybak (MPICE Jena) for advice regarding the numbers presented in **Figure 9C, D**; all errors in these numbers are ours.

---

**Figure S1.**
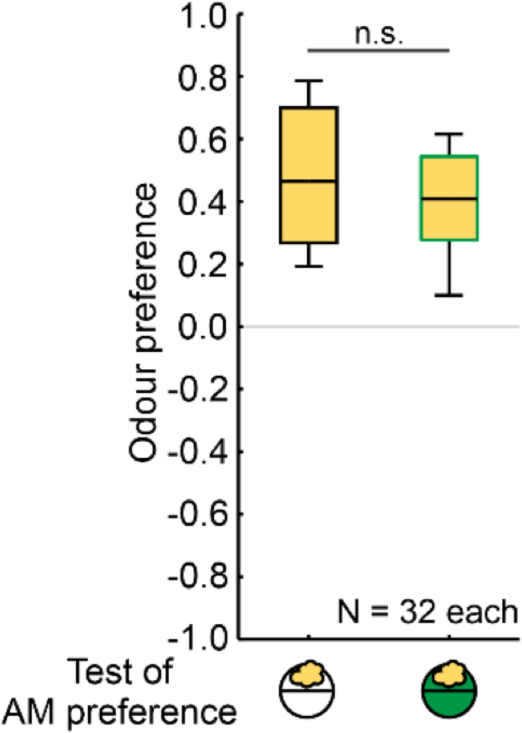
Naïve odour preference is unchanged in the presence of a sugar reward. Larvae were placed on a Petri dish filled with agarose (open circle) or sugar-supplemented agarose (green circle) together with one odour cup loaded with the indicated odour (yellow cloud) on one side and an empty odour cup on the other side of the Petri dish. After 3 min, the number of larvae on each side and in a middle zone were counted, and an odour preference was calculated according to Equation 1. The larvae showed approach to the odour regardless of whether the reward was present or not. Box plots represent the median as the midline, the 25/75% quantiles as box boundaries and 10/90% quantiles as whiskers. Sample sizes are indicated within the figure. n.s indicates non-significance in an MWU-test. Statistical results and source data in **Supplemental_Table_S1.xlsx**.

**Figure S2.**
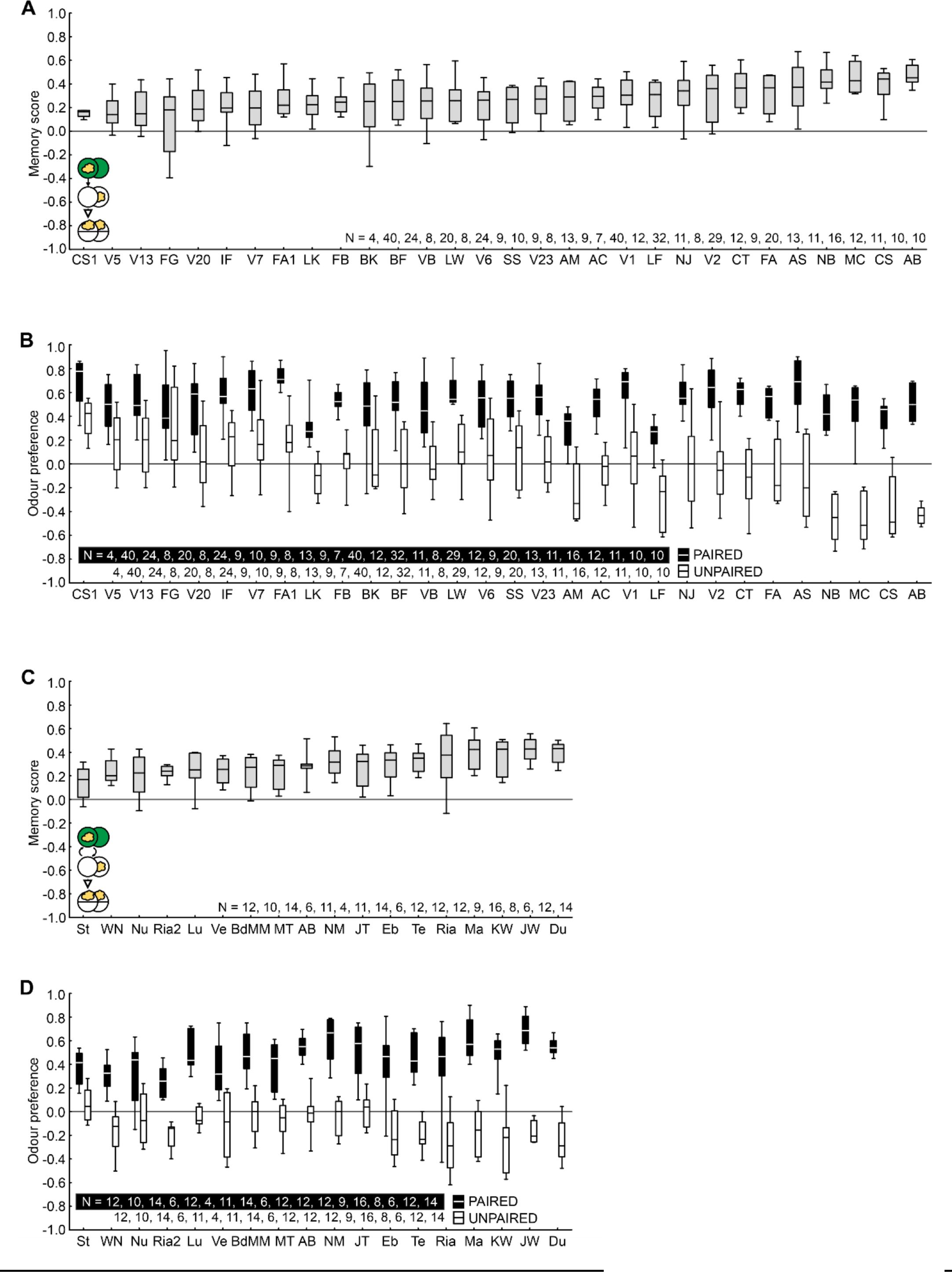
Separation of the pooled data depicted in Fig. 4. **(A)** Memory scores for the individual datasets underlying **Fig. 4B**, sorted by increasing median memory scores. **(B)** Odour preferences for the PAIRED (black boxes) and UNPAIRED (white boxes) trained groups underlying the memory scores in (A). **(C)** Memory scores for the individual datasets underlying **Fig. 4F**, sorted by increasing median memory scores. **(D)** Odour preferences for the PAIRED (black boxes) and UNPAIRED (white boxes) trained groups underlying the memory scores in (C). Letters below the plots refer to experimenters. Box plots represent the median as the midline, the 25/75% quantiles as box boundaries and 10/90% quantiles as whiskers. Sample sizes are indicated within the figure. Source data are given in the data file **Supplemental_Table_S1.xlsx**.

**Figure S3.**
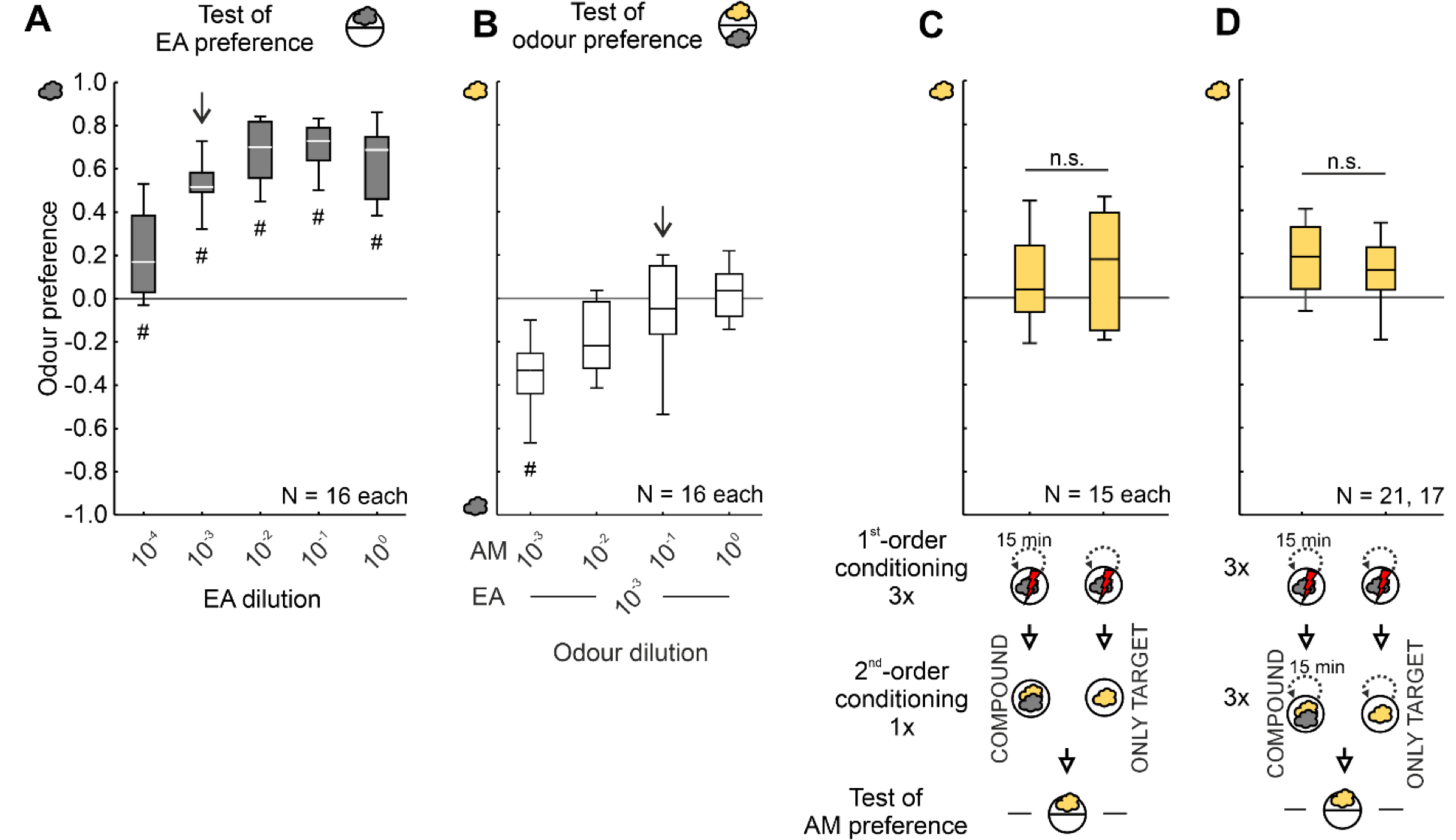
No evidence of second-order conditioning in the aversive domain. **(A-B)** Ethyl acetate (EA) (Sigma-Aldrich, Cat. No. 270989; CAS No. 141-78-6, diluted 10-3 in paraffin oil, CAS: 8042-47-5, AppliChem, Darmstadt, Germany) and amyl acetate (AM) (Sigma-Aldrich, Cat. No. 46022; CAS No. 628-63-7, diluted 10-1 in paraffin oil) were used as the trained and target odour, respectively. These dilutions were determined empirically to support moderate innate attraction to EA **(A)** (arrow) and a balanced choice between EA at such a moderately attractive dilution versus AM **(B)** (arrow). Paraffin oil is without behavioural significance to the larvae (Saumweber et al. 2011). Odour-electric shock associative learning experiments have previously been described in detail (Tomasiunaite et al. 2018). In brief, a custom-built semi-automatic device called “MaggotShock V 2.1” was used, which can transmit electric shock pulses through a custom-made setup with an 85-mm-diameter Petri dish filled with agarose (Sigma-Aldrich, Cat. No. A9539, CAS No. 9012-36-6). An electric shock stimulus from a DC power supply set to a total output of 80 V consisted of 30 pulses, each lasting 250 ms and followed by 250-ms breaks, for a total of 15 s. During conditioning, odours were presented on filter papers centred on the Petri dish lid. For the test, the larvae were transferred to the centre of a fresh 85-mm-diameter Petri dish equipped with a custom-made Teflon container with perforated lids and loaded with the target odour; these odour containers were randomly placed on the left or right side of the Petri dish. After 2 min the position of the larvae was noted, and a preference score for the target odour was calculated according to Equation 1. **(C-D)** During first-order conditioning, the larvae were placed onto the Petri dish, where they received the trained odour, followed 10 s later by electric shock for another 15 s. Coinciding with the offset of electric shock, the odour was removed, and the larvae were left undisturbed for 15 min. This cycle was repeated two more times. During second-order conditioning, the COMPOUND groups received the trained and the target odour in compound for 25 s, either only once (**C**) or three times (**D**, at 15-min intervals), whereas the ONLY TARGET group received the target odour only. For both groups, a test for the preference for the target odour followed. Lower odour preferences in the COMPOUND than in the ONLY TARGET group would offer preliminary evidence for second-order conditioning. No such difference was observed. Box plots represent the median as the midline, the 25/75% quantiles as box boundaries and 10/90% quantiles as whiskers. Sample sizes are indicated within the figure. # indicates significance from zero in an OSS test, n.s. indicates non-significance in an MWU-test. Source data are given in the data file **Supplemental_Table_S1.xlsx**.

